# Equilin does not affect thyroid hormone signaling in the developing *Xenopus laevis* tadpole brain

**DOI:** 10.1101/832071

**Authors:** Robert G. Bass, Zahabiya Husain, Lara Dahora, Christopher K. Thompson

**Affiliations:** School of Neuroscience, Virginia Tech, Blacksburg VA

## Abstract

Toxcast/Tox21 is a massive federally run research effort dedicated to better understanding the potential toxicity of thousands of compounds in a high throughput manner. Among this list of compounds is equilin, an estrogen-like compound that was flagged as a potential thyroid hormone agonist. Here we examine if equilin acts like a thyroid hormone agonist on cellular and molecular mechanisms of brain development in *Xenopus laevis* tadpoles. To examine the effect of equilin, tadpoles were divided into eight groups and received 4 days of exposure. The experimental groups were as follows: 1 μL, 10 μL, and 100 μL of equilin, 1 μL, 10μM, and 100 μM of 17-β estradiol as an estrogen control, 15 μg/mL thyroxine (T_4_) as a thyroid hormone control, and a no-exposure control. After 4 days of treatment, animals were treated with CldU to label dividing cells for 2hr and then euthanized in MS-222. After fixation, body length was measured and the brains dissected out. IHC was performed on brains for CldU to label proliferating neural progenitor cells. Brains were then whole-mounted and analyzed using confocal microscopy. We found that equilin did not increase the number of dividing progenitor cells in a T_4_-like manner. Instead, equilin decreased proliferation in a dose-dependent manner, as did estradiol. The same paradigm was performed separately staining for caspase-3 and h2ax, finding that equilin increased cell death in contrast to CNTL and T_4_. In another experiment, RNA was extracted from tadpole brains in each group and qPCR was performed to assess change in expression of thyroid hormone-sensitive genes, Equilin did not affect gene expression in a thyroid hormone-like manner. Our data indicate that equilin does not act as a thyroid hormone agonist in the *Xenopus laevis* nervous system but instead acts similarly to estradiol. Our data strongly suggest that equilin is not a TH disruptor, contrary to the findings of the ToxCast/Tox21 dataset.

## Introduction

Thyroid hormone (TH) is essential for normal brain development in vertebrates. Deficiencies in thyroid hormone, known as hypothyroidism, during development can result in significant and life-long mental and neurological deficits, including microcephaly(Rastogi & LaFranchi, 2010). Hyperthyroidism during pregnancy is less consequential but is associated with increased incidence of ADHD, autism, and epilepsy (S. L. Andersen, Andersen, Vestergaard, & Olsen, 2018; S. Andersen, Laurberg, Wu, & Olsen, 2014). Dysregulation of TH signaling can compromise neurogenesis, synaptogenesis, neuronal differentiation, and neuronal migration (Bernal, 2015; Howdeshell, 2002). There is considerable concern that endocrine disrupting chemicals (EDCs) that interfere with TH signaling may affect cognitive function via dysregulation of TH-dependent cellular and molecular mechanisms of brain development (Bernal, 2015; Colborn, vom Saal, & Soto, 1993; Lambert, Giller, Barber, Fitzgerald, & Skelly, 2015) Thus, substantial effort has been made to identify potential EDCs present in health and health products, chemicals used agriculture and industry, and others that make their way into the environment (Weiss et al., 2015).

The federally funded research program Toxicology in the 21^st^ Century (Tox21), has evaluated the potential for toxicity of thousands of compounds, including several tests for dysregulation of TH signaling. The GH3_TRE lucerferase cell assay in the Tox21 screening program identified equilin, a conjugated equine estrogen and primary ingredient of the drug Premarin,as a potential TH receptor (TR) agonist. Furthermore, previous studies have indicated that equilin, induces neurotrophism (Brinton, Proffitt, Tran, & Luu, 1997) and modulates circulating levels of the thyroid hormone thyroxine (T4) in humans. (Abdalla et al., 1987) In the present study, we test the hypothesis that equilin acts as a TH agonist in *Xenopus laevis* tadpoles, which are a widely used animal model for studies involving TH disruption toxicity. *Xenopus laevis* tadpoles are particularly sensitive to increases in TH signaling because it is the key initiator of metamorphosis (Sachs & Buchholz, 2017). We treated tadpoles 2-4 days of acute exposure to equilin and examined changes in body size, gene expression, and rates of neurogenesis and and cell death in the optic tectum, the functional homolog of the superior colliculus. We also compared the effects of equilin to tadpoles treated with 17β-estradiol, to serve as an estrogen control.

## Materials and Methods

### ANIMALS AND EUTHANASIA

We used albino *X. laevis* tadpoles that were bred on site. Tadpoles were reared in 0.1× Steinberg’s solution **(12 h light/dark cycle at 22°C)** and were selected for study based on their developmental stage using the morphological criteria of (Tarin, 1968) at the beginning of each experiment. *Xenopus* tadpoles free-swim 3–4 d after fertilization [stages 38–41, as defined by (Tarin, 1968). By 5–7 d, tadpoles have exhausted their yolk stores and begin to forage for food. We used tadpoles during these stages (stages 46–48) while the retinotectal circuit continues to undergo substantial development (Tao & Poo, 2005)(;(Bestman, Huang, Lee-Osbourne, Cheung, & Cline, 2015; Bestman, Lee-Osbourne, & Cline, 2012a);(Deeg, Sears, & Aizenman, 2009) (Pratt & Aizenman, 2009). Under standard rearing conditions in our laboratory, stage 46 lasts 1 d, stage 47 continues for 2–3 d, stage 48 lasts 5 or more days. During treatment bath experiments, we kept the tadpoles in bowls with 200 ml of 0.1× Steinberg’s solution (see below for details about hormone experiments). Tadpoles were fed tadpole powder (Xenopus Express) once a day, and waste was removed as needed. Euthanasia was performed via treatment with 0.2% MS222 until the heart was stopped. All animal procedures were performed in accordance with Virginia Tech’s institutional animal care and use committee’s regulations. The number of animals used in each experiment is listed in the figure legends.

### HORMONE TREATMENTS

For T_4_ treated tadpoles, 100mg T_4_ (Sigma) was dissolved in 6.66mL of 50nM NaOH to form a 1.93mM stock solution stored at −20°C. Working stock solution was made by diluting original stock down to 1.93μM in Steinburg’s solution. To make 15 μg/L T_4_, 2mL of working stock was added to 200mL of Steinberg’s solution tadpole bath along with 200 μ DMSO. For equilin treated tadpoles, 100mg equilin (Sigma) was dissolved in 3.72 mL DMSO to form a 0.1M stock solution stored at −20°C. For 100 uM, 10 uM, and 1 uM equilin-treated groups, the following volumes of stock were introduced respectively into 200 mL tadpoles baths: 200μL stock, 20 μL stock with 180 μL DMSO, 2 μL stock with 198 μL DMSO. For 17β-estradiol treated tadpoles, 300mg estradiol (Sigma) was dissolved in 11.01 mL DMSO to form a 0.1M stock solution stored at −20°C. For 100 μM, 10 μM, and 1 μM estradiol-treated groups, the following volumes of stock were introduced respectively into 200 mL tadpoles baths: 200μL stock, 20 μL stock with 180 μL DMSO, 2 μL stock with 198 μL DMSO.

### CHLORO-DEOXYURIDINE INCORPORATION TO LABEL DIVIDING CELLS

Chloro-deoxyuridine (CldU; MP Biomedicals) was dissolved in distilled H_2_O to generate 38.1 mM stock, which was then diluted to 10 mM in Steinberg’s solution. We placed tadpoles in 10 mM CldU solution for 2 h and then euthanized them immediately. Tadpoles were then fixed in 4% PFA until needed for immunostaining.

### WHOLE-MOUNT IMMUNOSTAINING AND IMAGING. (DIVIDING CELLS)

Tadpole brains were washed in PBS-TX, placed in blocking buffer (2.5% normal goat serum in PBS-TX) for 1 h, and incubated overnight in the primary antibody [5-chloro-2’-deoxyuridine (CldU), 1:1000, NOVUS biologicals, NB500-169, made in rat; in blocking buffer at 4°C with gentle rotation. We then washed the brains three times in PBS-TX, incubated them in the appropriate secondary antibody [CldU with Alexa Fluor 488 (1:500), Abcam] for 3–4 h, washed them again in PBS-TX, and incubated them for 15 min in Sytox-O (1:500 in PBS). Brains were washed in PBS once and cover-slipped in a well on a slide with mounting medium composed of 3.6 g urea, 5 mL of glycerol, and 5 mL of Millipore purified water. Immuno-stained brains were imaged on a Leica SP8 confocal microscope.

### WHOLE-MOUNT IMMUNOSTAINING AND IMAGING. (DYING CELLS)

Tadpole brains were washed in PBS-TX, placed in blocking buffer (2.5% normal goat serum in PBS-TX) for 1 h, and incubated overnight in the primary antibody (h2ax, 1:500, EMD Millipore, 05-636, made in mouse; and caspase 3, 1:500, abcam, AB13847, made in rabbit; in blocking buffer at 4°C with gentle rotation. We then washed the brains three times in PBS-TX, incubated them in the appropriate secondary antibody [h2ax with Alexa Fluor 488, 1:400, Life Technologies, A11029, goat anti mouse; caspase-3 with Alexa Fluor 647, 1:400, Invitrogen/Thermofisher scientific, A21244, goat anti rabbit] for 3–4 h, washed them again in PBS-TX, and incubated them for 15 min in Sytox-O (1:500 in PBS). Brains were washed in PBS once and cover-slipped in a well on a slide with mounting medium. Immuno-stained brains were imaged on a Leica SP8 confocal microscope.

### QUANTIFICATION OF CELL NUMBER

We counted the number of immuno-positive cells from optical stacks of whole imaged tecta in Imaris. A rendering of 70 transversal cross-sections within the optic tectum was rendered in 3 dimensions and the density of dying cells was analyzed using. The area of the tectum that was quantified was delineated by the anterior commissure on the rostral side, the most caudal extent of the midbrain ventricle, the dorsal surface of the brain, and the top 70 optical sections of tectum were used to delineate the ventral extent of the region of interest. For CldU, we counted cells that had an intensity of fluorescence at least 2X background.

### QUANTIFICATION OF OPTIC TECTUM SIZE

The same blinded microscope images from the cell death experiments were used to quantify changes in overall brain volume in the optic tectum. For each image, the 70 transversal cross sections in the file were projected into one image. The region of interest was delineated by the anterior commissure on the rostral side, the most caudal extent of the midbrain ventricle, the dorsal surface of the brain, and the top 70 optical sections of tectum were used to delineate the ventral extent of the region of interest. Optic tectum size was calculated in ImageJ using the polygon sectioning tool to measure the area of the selected region of interest.

### QUANTIFICATION OF GENE EXPRESSION

We performed two experiments that quantified changes in gene expression. First, eight groups of 18 stage 46 tadpoles were exposed to treatments in the following groups: 15 ug/mL T4, Equilin 1 μM, 10 μM, 100 μM, 17-β Estradiol 1 μM, 10 μM, 100 μM, and a no exposure control. The exposure lasted 4 days, after which the animals were euthanized by an overdose of MS222 (0.2%). We quickly dissected the brains rostral to the hindbrain–spinal cord junction, placed them into Trizol (Life Technologies), and froze them in −80°C. We extracted RNA per the manufacturer’s instructions for Trizol, measured the amount of RNA extracted on a NanoDrop, and reverse transcribed the mRNA using the iScript kit (Bio-Rad), using 1000 ng of RNA per reaction. We performed quantitative PCR (qPCR) using 500 ng of cDNA per reaction using the iTaq Universal SYBR Green Supermix kit (Bio-Rad) on a Bio-Rad CFX384 thermocycler. A description of primers used is in <u>Table 1</u>. All reactions were done with technical triplicates; outlier reactions (deviations more than 0.5 times the SD from the mean) within a set of triplicates were removed from analysis. We used a two-step reaction with a 2 min 95° melt step, followed by a 30 s 60° annealing and an extension step for 40 cycles, with fluorescence measured at the end of every 60° step. At the end of 40 cycles, we evaluated the melt curves for secondary products. None of the primers used in this study generated secondary products. Four genes were utilized which were previously identified as sensitive to TH signaling. (Das et al., 2006; Denver, Pavgi, & Shi, 1997) the selected genes were as follows: nrep (For: TGTAGCGGGAGCAATCACAA, Rev: ACATTCAGAAACCTGCCCCT, amplicon size 162) pcna (For: TTCTTGTGCGAAGGATGGGG, Rev: CGCAATGCAAATGTGAGCTG, amplicon size: 150), cycs (For: TCAAATCAAGGTGCAGTCAT Rev: TTAGGTCCAGTTTTGTGCTT amplicon size:) and thrβ (For:AGATCATGTCCCTCCGAGCA, Rev: CCACACCGAGTCCTCCATTT amplicon size: 114)

We used two genes for use as a reference as outlined in (Thompson & Cline, 2016): *rps13* (For; ATGTCAAGGAACAGATCTTCAAACT, Rev: GAGGATTCTCAGGATTTTATTACCA, amplicon size: 131) and *rpl32* (For: GCTCTTCATCGGGCTGTCTA, Rev: CTTGGTGAGAGGTCTGAGGG amplicon size: 77)(we evaluated the expression of target genes using the comparative Ct method against the mean expression per brain of these genes.

### STATISTICS

We used GraphPad Prism to analyze our data for statistical differences. We used regression analysis to evaluate the relationship of optic tectum volume. Data with only two groups were evaluated with Student’s *t* test. Data sets with more than two groups were evaluated with ANOVA, and *post hoc* comparisons were made with Tukey’s multiple comparisons test. Data sets with two dimensions (e.g., time vs treatment) were evaluated with two-way ANOVA, and *post hoc* comparisons were made with either Tukey’s multiple comparisons test. We set α = 0.05 for all tests.

## Results

### Equilin exposure caused distinct craniofacial morphology change

Previous data show that thyroid hormone treatment causes distinct morphology changes in head shape. (Thompson & Cline, 2016) To assess if similar craniofacial changes were induced by equilin, tadpoles were placed in one of the following exposure groups for 4 d: 15 ug/mL of T_4_, 100 μM equilin, 100 μM 17β-estradiol, or no-exposure control (Steinburg’s rearing solution). Tadpoles were then euthanized in MS-222 and images were taken from various angles and are depicted in Figure 1A. Equilin-associated morphology change appeared to require a shorter onset period than that of thyroid hormone. Equilin induced substantial changes in craniofacial morphology, but many of these changes differed from what is seen in thyroid hormone-treated tadpoles. Most notable was an abrupt narrowing of the mouth and vertical orientation of the lower jaw. There is some extension of Meckle’s cartilage, similar to thyroid hormone, but narrowing of the mouth is more pronounced. There were no visible changes in gut length. The snout is significantly smaller and directed more dorsally than what is typically seen in TH-induced metamorphic-like changes in head anatomy. The expression of this phenotype was inconsistent; nearly half of equilin-treated animals produced a normal facial morphology (such as in Fig2). Given the These data indicate that equilin is at the very least acting on non-TR mechanisms perhaps in addition to TR-mediated mechanisms.

**Figure 1).**
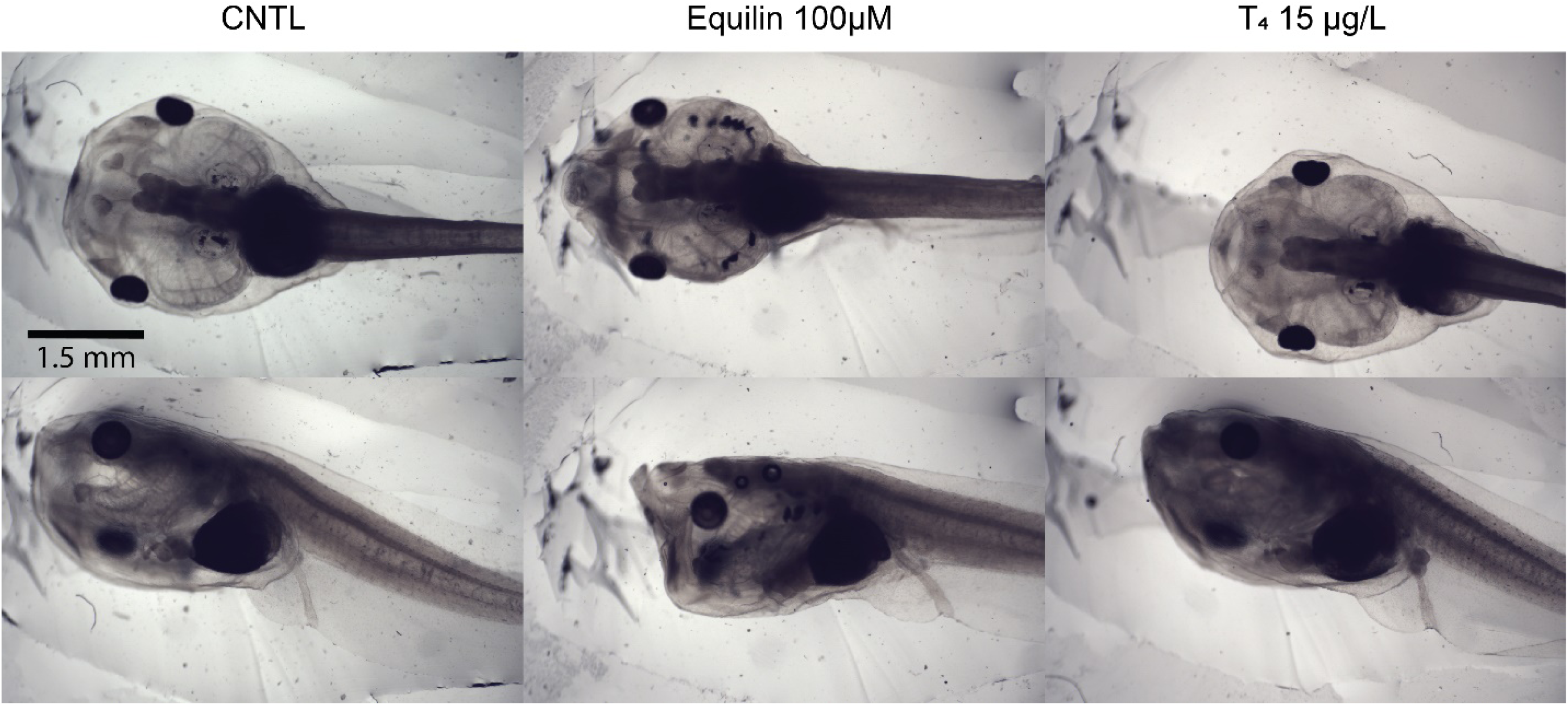
Equilin-induced craniofacial morphology change does not resemble T_4_. Photomicrographs of the dorsal (top row) and lateral (bottom row) aspects of sibling tadpoles in control (CNTL, left) equilin (middle), and T_4_ (right) treatments for four days. Equilin induced substantial changes in craniofacial morphology, but many of these changes differed from typical changes seen in thyroid hormone-treated tadpoles. Most notable was an abrupt narrowing of the mouth and vertical orientation of the lower jaw. There was some extension of Meckle’s cartilage, similar to thyroid hormone, but narrowing of the mouth was more pronounced. There were no visible changes in gut length.

### Equilin decreased cell proliferation in the optic tectum

To determine if equlin treatment induces thyroid hormone-like changes in mechanisms of brain development, we examined features of the optic tectum, a region of the midbrain. The retino-tectal circuit is a simple and well-characterized component of the amphibian brain commonly used to study neuro-development. (Liu, Hamodi, & Pratt, 2016) Neural progenitor cells (NPCs) in the optic tectum are located adjacent to the midbrain ventricle. NPCs give rise to neurons that are subsequently displaced laterally from the midbrain and become the neuronal cell body layers (Bestman, Lee-Osbourne, & Cline, 2012b). To test the hypothesis that equilin exposure induces thyroid hormone-like changes in tectal cell proliferation, we used treated tadpoles with CldU 2 hr prior to sacrifice and used immunohistochemistry (IHC) to label proliferating cells within the optic tectum. Density of the CldU+ cells was calculated based off the volume of the midline of each brain. T_4_ increased cell proliferation relative to CNTL as anticipated (Fig 2, p < 0.0001, one-way ANOVA, Tukey’s multiple comparison test). In contrast, 17-β estradiol and equilin induced significant decreases in cell proliferation relative to CNTL (Figure 2, p < 0.0001, one-way ANOVA, Tukey’s multiple comparison test). Thus, equilin induces an estrogen-like effect and not a thyroid hormone-like effect on neuronal proliferation in the developing brain.

**Figure 2).**
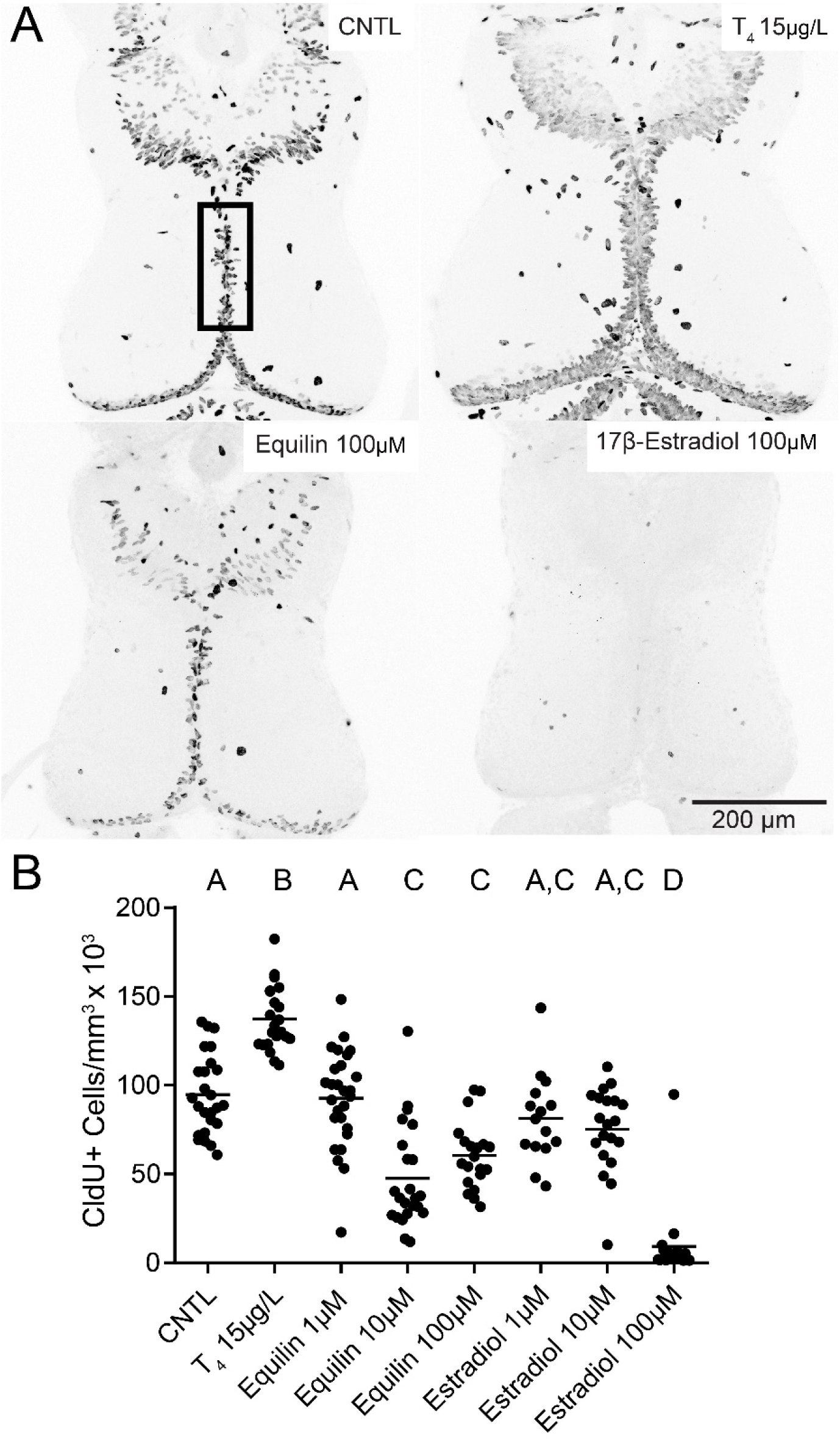
Equilin and estradiol reduces cell proliferation in the optic tectum. A) z-projected images of tadpole midbrains from four of eight treatments immuno-stained for CldU to identify proliferation cells at sacrifice. The greyscale has been reversed from the original images for presentation purposes. B) Scatter dot plot of density of CldU+ proliferating cells along the midline. T_4_ significantly increased proliferation in the optic tectum, whereas equilin and estradiol decreased proliferation in a dose dependent manner relative to CNTL. Letters above groups designate post-hoc significant differences.

### Equilin increased cell death in optic tectum

The estrogen and equilin-treated brains from the CldU experiment appeared to be smaller than CNTL, suggesting that these treatments may increase cell death in the brain. To test this hypothesis, we treated tadpoles with 15 ug/mL T_4_, Equilin 100 μM, 17-β Estradiol 100 μM, or CNTL for four days. Brains were then immuno-stained for h2ax and caspase-3 to stain for dying cells and Sytox-O as a general cell marker but also was used to identify dying cells (Faulkner, Wishard, Thompson, Liu, & Cline, 2014; Thompson & Cline, 2016). Representative z-projection images were taken on a confocal microscope. Z-projections were then rendered in ImageJ and cells were counted using the designated cell-counting ImageJ extension. For h2ax and caspase-3, cells with increased fluorescence relative to other cells were designated as dying and were thus counted. Additionally, fluorescent cells in the cell body layer adjacent to the midbrain ventricle were counted for sytox-O. (Figure 3) Equilin treatment significantly increased cell death relative to control in all three markers for cell death (Figure 3, p < 0.0008 for caspase-3, p<0.0001 for h2ax, p<0.0001 sytox-O, one-way ANOVA, Tukey’s multiple comparison test) whereas T_4,_ had no effect, and trended towards a decrease in cell death. (Figure 3, p < 0.7745 for caspase-3, p<0.0982 for h2ax, p<0.9900 sytox, one-way ANOVA, Tukey’s multiple comparison test). Estradiol increased cell death as well, albeit only in intensely-stained Sytox-O-stained cells. (p < 0.0001, one-way ANOVA, Tukey’s multiple comparison test) This may be due in part to the fact that Sytox-O+ dying cells tends to be a late-stage cell death marker, and that increases in the early stage markers like H2AX and caspase-3 may have occurred prior to sacrifice.

**Figure 3).**
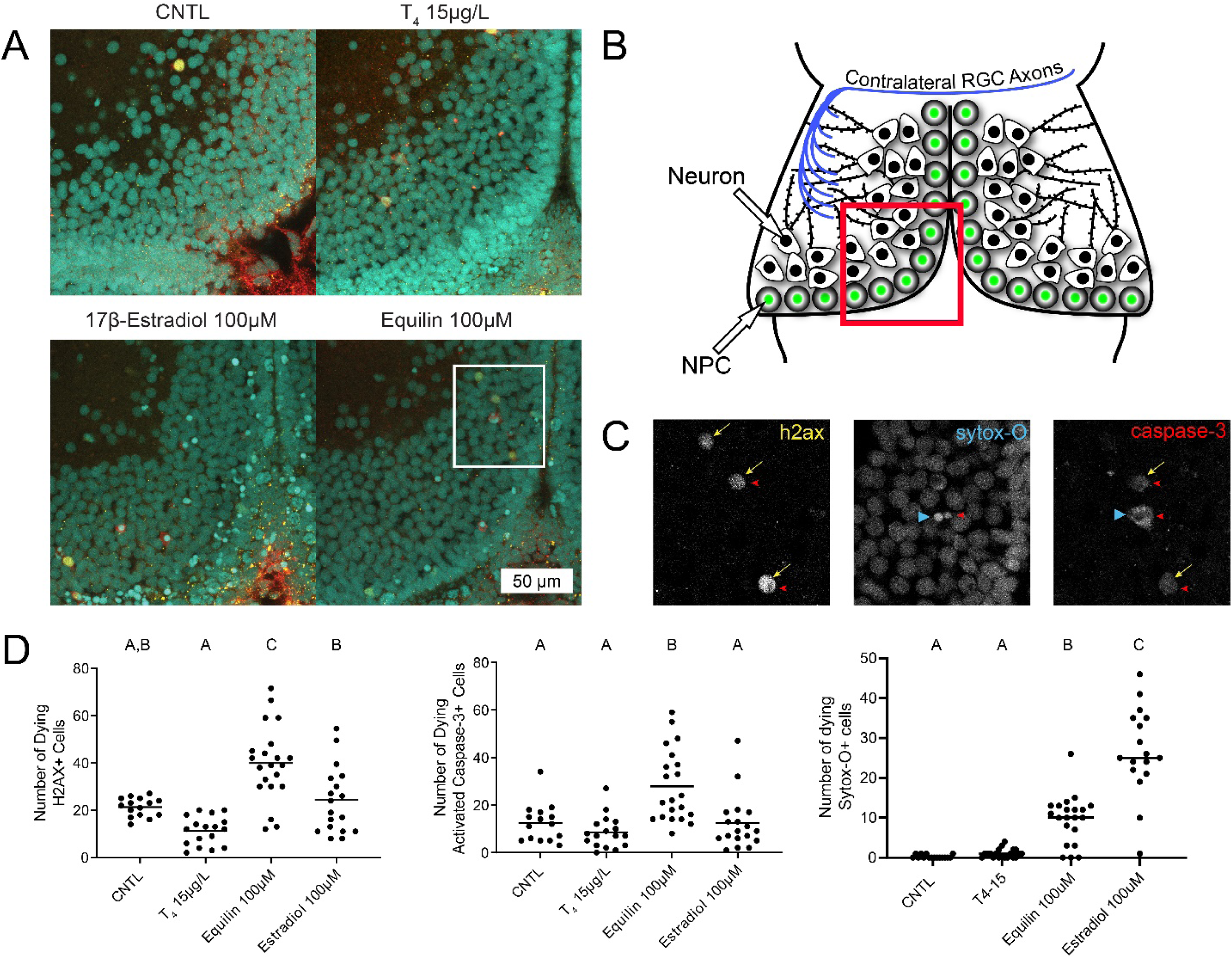
Equilin increased programmed cell death. A) Z-projections of a portion of optic tectum from example brains in each group. IHC was performed for H2AX (yellow) and activated caspase-3 (red), and brains were counterstained for Sytox-O (teal). B) Diagram of the optic tectum illustrating the location of example images in A. C) Separate channels of the three stains from the inset in the equilin-treated example. Arrows designate h2ax+ cells, whereas arrowheads designate both caspase-3+ and dying Sytox-O+ cells, which are much brighter than healthy cells. D) Equilin increased the number of dying cells relative to CNTL in all three measurements. Estradiol significantly increased the number of dying Sytox-O+ cells. Letters above groups designate post-hoc significant differences.

### Equilin failed to increase expression of thyroid hormone-sensitive genes

Thyroid hormone influences the expression of hundreds of genes in the developing *Xenopus brain.* (Das et al., 2006; Denver et al., 1997) We used RT-PCR to determine if equilin alters expression of thyroid hormone-sensitive genes similar to T_4_. Tadpoles were treated with 15 μg/L T_4_, three concentrations of equilin, three concentrations of estradiol, or CNTL for 4 days, mRNA was extracted from brains, which was reversed-transcribed and analyzed via RT-PCR. Thyroid hormone-sensitive genes were upregulated in the thyroid-hormone treated brains, as expected. (Figure 4, p<0.0001 for NREP, p<0.0001 for PCNA, p< 0.0001 for cycs, p<0.0031 for thrβ, one-way ANOVA, Tukey’s multiple comparison test) Equilin treatment, on the other hand, failed to produce the same increase in gene expression and did not differ from CNTL (Fig 5, p<0.9997 for NREP, p<0.0.9927 for PCNA, p< 0.0.4932 for cycs, p<0.9908 for thrβ, one-way ANOVA, Tukey’s multiple comparison test). These results strongly suggest that equilin does not activate thyroid-hormone sensitive genes in the developing tadpole brain.

**Figure 4).**
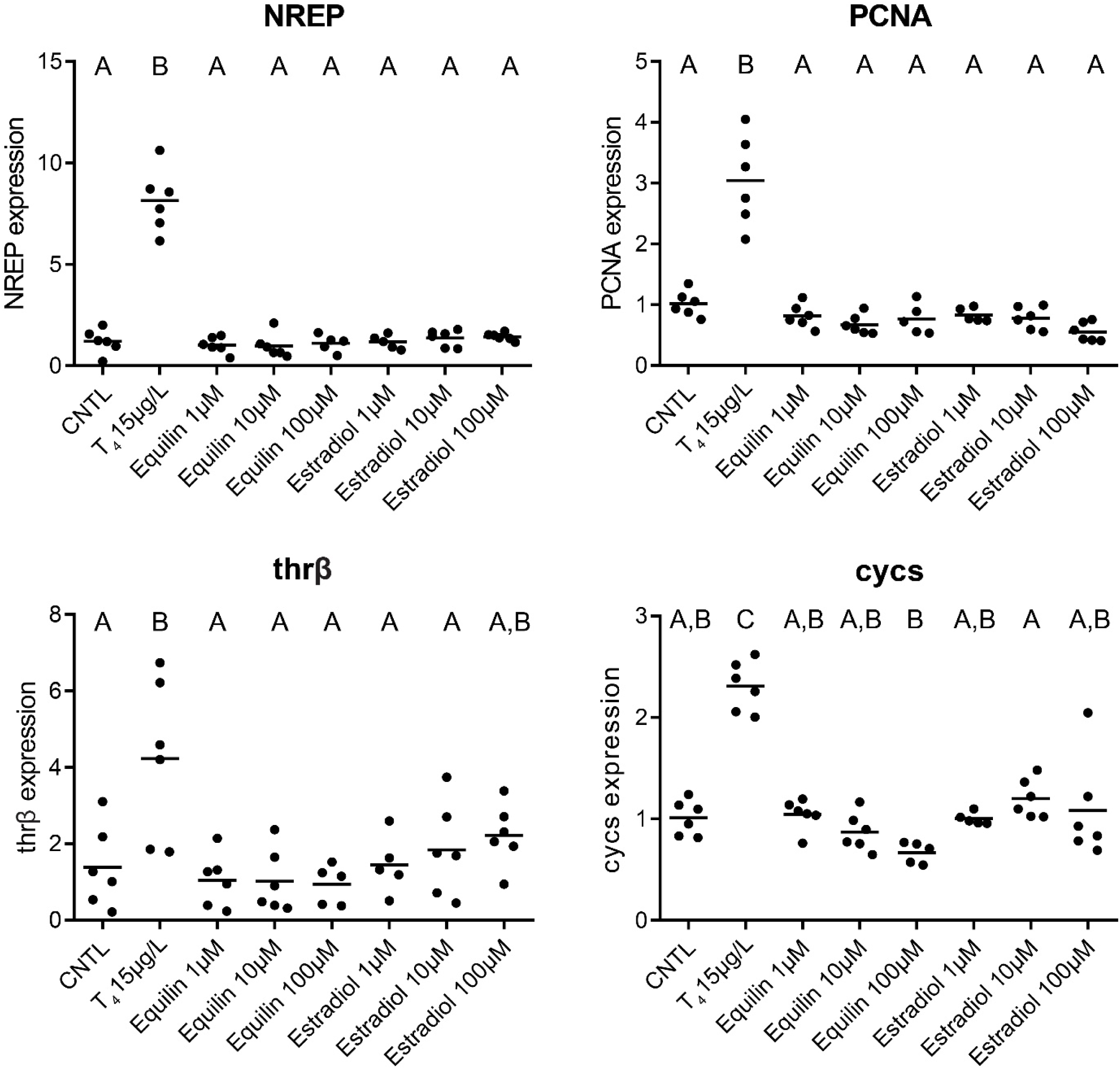
Equilin did affect expression of thyroid hormone-sensitive genes. Thyroid hormone treatment significantly increased expression of neuronal regeneration related protein (NREP), proliferation cell nuclear antigen (PCNA), thyroid hormone receptor beta (thrβ), or somatic cytochrome c (cycs) relative to control. Equilin and estradiol had no effect on expression of these TH-sensitive genes. Letters above groups designate post-hoc significant differences.

## Discussion

Our data provide no support for the hypothesis that equilin functions as a thyroid hormone agonist. Equilin fails to replicate the same genetic and morphological changes of the brain seen in animals exposed to thyroid hormone. In fact, in some instances, it produces the opposite effect, such as a decrease in NPC proliferation (Fig 3), which stands in contrast to the effects of thyroid hormone exposure on proliferation (Thompson & Cline, 2016). These observed effects of equilin -exposed animals suggest similarity to that of estradiol exposed animals, indicating that equilin is functioning in this region as an estrogen receptor (ER) agonist.

### Equilin is an ER agonist

Estradiol is an important regulator brain development in vertebrates, (Beyer, 1999; Coumailleau et al., 2015). It contributes to brain plasticity, cell migration, cell proliferation, dendritic spine growth(Bhavnani & Stanczyk, 2014; Diotel et al., 2013). These effects are conserved across many vertebrates; However, they only appear late in development or adulthood. Exposure to estradiol during embryonic or fetal development typically retards development and can be lethal. In *Xenopus laevis*, it has been shown to severely disrupt metamorphosis, ER gene expression, body morphology, and promote lethality (Nishimura, Fukazawa, Uchiyama, & Iguchi, 1997) in stages 1-39 of development. These developmental stages are analogous to early embryonic stages of development in mammals. Our results are consistent with these previous reports, in that estradiol induced similar effects on the head and promoted cell death in the brain, which would lead to lethal conditions. The equilin-exposed groups were very similar to these reported effects of estradiol, which provides strong evidence that equilin acts like an estrogen compound in developing tadpoles. This makes sense give that equilin, when used as a pharmaceutical ingredient, serves as one of many conjugated equine estrogens that compose estrogen-therapies such as *Premarin*. It functions as a partial ER agonist, with higher binding affinity for the ERβ receptor subtype rather than ERα (Bhavnani & Stanczyk, 2014). The binding affinity of equilin to both of these nuclear ERs is lower than that 17β-estradiol, which may explain why 17β-estradiol was more effective on reducing cell proliferation than equilin at lower doses (Fig 3).

### Equilin does not act like a TR agonist in the developing brain

The effects of equilin stand in stark contrast to the effects of thyroid hormone on brain development in tadpoles, including increases in neurogenesis, brain volume, and change in expression of thyroid hormone-sensitive genes (Thompson & Cline, 2016). These changes are facilitated largely by action on TR, primarily the TRα subtype. There is considerable evidence of crosstalk between ERs and TRs. However, these reviews primarily discuss the potential action of thyroid hormone on ER, providing no evidence that ER agonists such as equilin may have crosstalk with TR (Hogan, Crump, Duarte, Lean, & Trudeau, 2007; Vasudevan, Ogawa, & Pfaff, 2002). Nevertheless, our results provide no genetic or morphological evidence for that hypothesis. It therefore is clear that equilin does not act like a thyroid hormone agonist in the developing tadpole brain, which calls into question the results pertaining to equilin in the Tox21 project.

